# Transformer Graph Variational Autoencoder for Generative Molecular Design

**DOI:** 10.1101/2024.07.22.604603

**Authors:** Trieu Nguyen, Aleksandra Karolak

## Abstract

In the field of drug discovery, the generation of new molecules with desirable properties remains a critical challenge. Traditional methods often rely on SMILES (Simplified Molecular Input Line Entry System) representations for molecular input data, which can limit the diversity and novelty of generated molecules. To address this, we present the Transformer Graph Variational Autoencoder (TGVAE), an innovative AI model that employs molecular graphs as input data, thus captures the complex structural relationships within molecules more effectively than string models. To enhance molecular generation capabilities, TGVAE combines a Transformer, Graph Neural Network (GNN), and Variational Autoencoder (VAE). Additionally, we address common issues like over-smoothing in training GNNs and posterior collapse in VAE to ensure robust training and improve the generation of chemically valid and diverse molecular structures. Our results demonstrate that TGVAE outperforms existing approaches, generating a larger collection of diverse molecules and discovering structures that were previously unexplored. This advancement not only brings more possibilities for drug discovery but also sets a new level for the use of AI in molecular generation.

## INTRODUCTION

The domain of medicinal chemistry continues to face exploratory phase, directing researchers to investigate the new chemical spaces for novel drug discovery. Particularly with the emergence of drug resistance, increased healthcare costs, and the continuous challenge of adverse side effects associated with current treatments, there is an urgent need to discover novel therapeutic agents. This urgency is particularly evident in the context of diseases as cancer, often lacking effective treatments or developing resistance mechanisms. Artificial intelligence (AI) has emerged as a transformative tool in the field of medicinal chemistry, offering potential to accelerate the drug discovery process. By employing the power of machine/deep learning (ML/DL) and neural networks (NN) algorithms, or natural language processing (NLP) models, AI can analyze vast repositories of chemical and biological data, recognize complex patterns, predict molecular interactions and optimize drug properties with high precision. The ability of AI algorithms to recognize hidden patterns and relationships within data makes it an invaluable tool in the design of novel, comprehensive, and targeted drug candidates from scratch or from existing libraries. Consequently, AI-enabled design of novel drug libraries can vastly increase the effectiveness of drug discovery processes, since the process starts with already drug-like molecules or potent ligands.

As we present in this article, AI can help explore the vast chemical space to generate novel compounds that meet specific medicinal chemistry criteria. Current models can predict the biological activity, pharmacokinetics, and toxicity of these newly designed molecules, guiding the optimization process to enhance their drug-like properties^1^. Additionally, AI-driven generative models, can propose innovative molecular structures that might not be discovered through traditional methods^2-5^. This capability to design unique and highly specific drugs holds great promise for addressing unmet medical needs and confronting diseases with limited treatment options. Unlike traditional drug discovery, which often relies on modifying existing compounds, generative molecular design allows scientists to create new molecules tailored to specific biological targets. This approach can lead to the development of drugs with enhanced specificity, reduced side effects, and improved efficacy. Moreover, generative molecular design is particularly valuable for addressing diseases with no current treatments or those resistant to existing therapies^6,7^. In addition to accelerating the identification of promising drug candidates, AI can also reduce the costs and time associated with traditional drug discovery methods. However, despite the promises of AI-based drug design, challenges and limitations persist. One recognized problem is limited resources of the high-quality, annotated datasets, which are essential for training the AI models. Furthermore, the intrinsic complexity of biological systems remains a challenge, because AI models can only capture the scale of molecular interactions as accurately as the molecular representation they see.

In the last decade, many AI-driven models for molecular generation and design have emerged. These models often use the powerful and widely recognized Simplified Molecular Input Line Entry System (SMILES)^8^ or other string representations of molecules. Alternatively, graph representations where nodes correspond to atoms and edges to bonds were applied. These approaches included models capable of generation of the new chemical compounds, e.g., CharRNN^9^, which implemented using Long Short-Term Memory^10^ network or LatentGAN^11^ which then use Generative Adversarial Network^12^ to train and learn from SMILES, to name a few. However, there are concerns associated with these models. One major concern is the ambiguity in representation, as a single molecule can be represented by multiple valid SMILES strings. Additionally, while SMILES can denote stereochemistry through various symbols, these additions increase the complexity of the data. These factors complicate the learning process for models that rely on string as input. In addition, there is a risk of algorithmic bias and overfitting associated with these models, which can compromise the reliability and generalizability of AI-generated predictions.

We address a number of these challenges by development of the Transformer Graph Variational Autoencoder (TGVAE) model for molecular design. The model emerges from the existing platform for molecular generation models called Molecular Sets (MOSES)^13^. The objective of TGVAE is to address the constraints of the string representation, which can be insufficient in capturing the complexity of molecular structures. Our technique utilizes Graph Neural Network^14,15^ (GNN) to train on molecular graph data, resulting in a more complete and accurate representation of chemical structures.

## DATASETS and METHODS

### Datasets

We utilized the MOSES^13^ dataset in our training and evaluation processes, aiming to compare our results with their baseline SMILES string-based model. The MOSES dataset is derived from the ZINC Clean Leads collection^16^, which has been filtered to include molecules with a molecular (MW) ranging from 250 to 350 g/mol, a maximum of seven rotatable bonds, and an logP^17^ value not exceeding 3.5. Molecules in the dataset include C, N, S, O, F, Cl, Br, H atoms, and the molecules with cyclic rings larger than eight atoms and molecules with charged atoms are removed following MOSES approach^13^.

For validation purposes we utilize the MOSES dataset parts: total consisting of 1,936,962 molecules with MOSES original split ratio for the training set = 1,584,664 molecules, the test set = 176,075 molecules, and the scaffold test set (TestSF) = 176,226 molecules. All molecules in the TestSF contain Bemis-Murcko scaffolds^18^ that are not found in either the training or test sets. This set serves as a critical evaluation measure to determine whether a model can generate novel structures.

### Data Representation

Molecules in the dataset are represented in SMILES format. SMILES strings are converted into three matrices to be processed by GNN (**Figure 1A**):

*Node features matrix V*_(1×*n*)_: Each column of this matrix is a feature of each node (atom) in the molecule. To align with other string-based models that only use character token as features, we only extract one feature for each node which is the atom’s symbol.

*Edge index matrix* 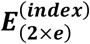: This 2D matrix illustrates how atoms in molecules are connected. Each column represents a bond connection, with the upper value indicating the starting atom and the lower value representing the ending atom. Columns are reversed to represent connections in both directions.

*Edge feature matrix* 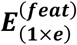: This matrix denotes bond types associated with each column of the edge index matrix. Bond types include single (0), double (1), triple (2), and aromatic (3).

### Model Architecture

The model builds upon the foundation of Transformer^19^, adapted for molecular design. It is an encoder-decoder architecture, in which the encoder learns properties from a molecular graph and encodes them into a latent space. The decoder then samples from this latent space to generate new molecules. Before entering the encoder, SMILES strings are converted to graph data (for details see the previous section on *Data Representation*). Node features, edge indices, and edge features are then inputted to Graph Attention Network^20^ (GAT) to embed node features to higher dimension. GAT is improved to use an attention mechanism to selectively learn information from nodes’ neighbors. This allows atoms in molecules to focus on important and highly related neighbor molecules and disregard irrelevant ones, which improves feature learning. The encoder is stacked with *N* encoder layers. Within every layer, we replace the self-attention layer in the original framework with two GAT layers. Subsequently, node features are directed to a fully connected feed-forward network. Residual connection^21^ followed by layer normalization is employed for all 3 sublayers. The model architecture is illustrated in **Figure 1B**. For model implementation we utilized Python programming language^22^ and PyTorch deep learning library^23^.

**Figure 1.**
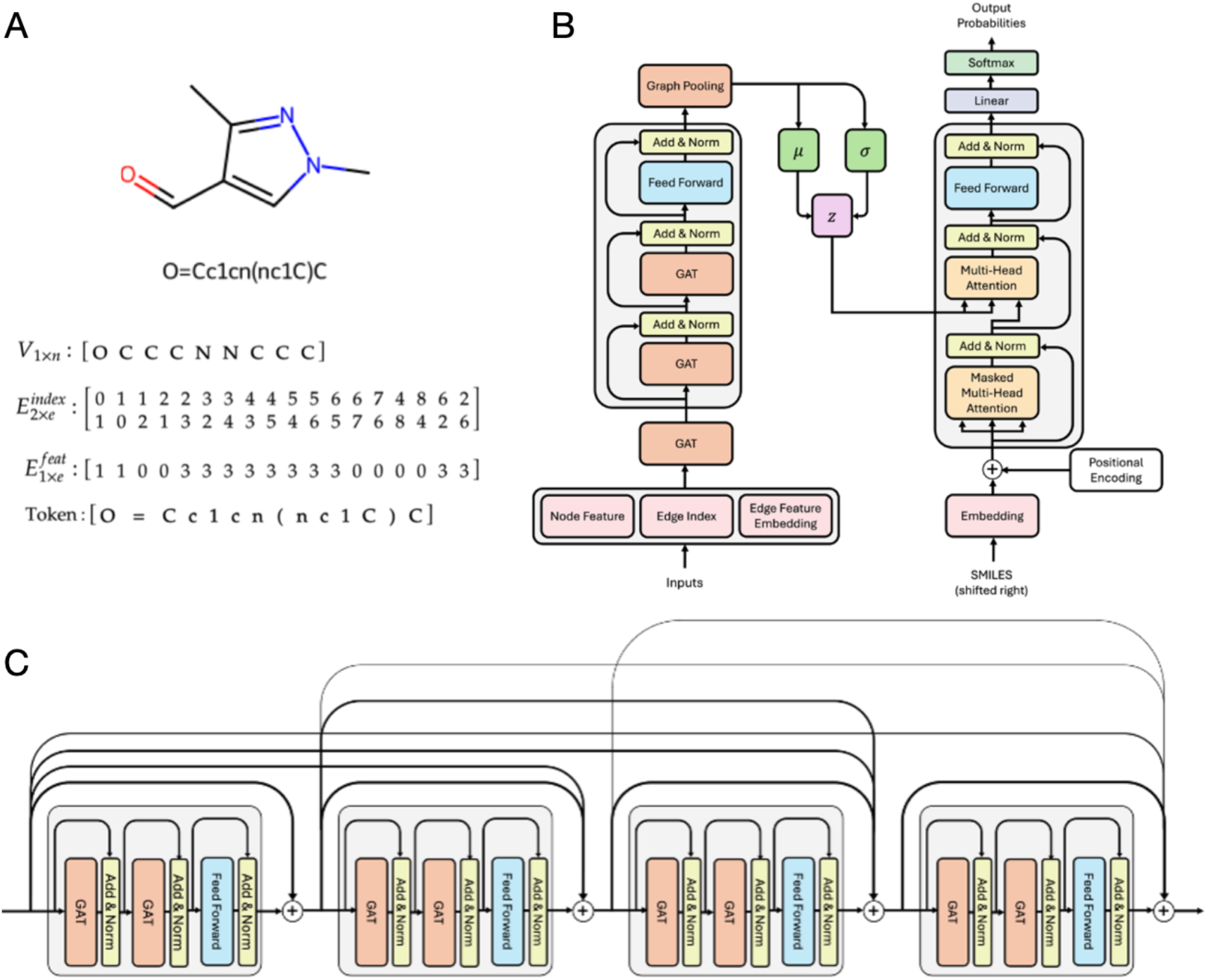
Molecular representation used in the TGVAE model. (A) Molecule structure and SMILES string with vectors definitions. (B) Architecture of the TGVAE model. (C) Connection of layers in TGVAE.

A stack of *N* encoder layers results in a deep model, which potentially can cause over-smoothing, a common issue seen in training deep GNN models. Over-smoothing refers to a phenomenon that representation of nodes in a graph become increasingly similar or indistinguishable. As a result, deep GNNs see a decrease in their ability to differentiate and classify data. It has been shown that ResNet^21^ and DenseNet^24^ are capable of overcoming this issue^25^. We overcome this issue by employing another dense connection for each encoder layer as shown in **Figure 1C**. This connection helps with vanishing gradients as each layer has a direct path to the final loss calculation. It also enhances feature propagation, where features learned in the earlier layers can be readily accessible by the later ones. These benefits have been shown to stabilize model’s training and significantly improve its performance. Following the last encoder’s layer, a readout layer called global add pooling is applied to aggregates of all nodes’ features to produce graph-level outputs. In the standard Transformer architecture, the encoder generates a contextual representation or learned memory from the input, which is processed and attended to by the decoder to generate outputs. TGVAE passes this learned memory through two linear layers (*μ* and *σ*) to convert it to a latent space distribution. The decoder retains its original structure, yet instead of attending to the memory from the encoder, the attention layer attends to the latent space. Utilizing information learned from molecular graphs, the decoder generates SMILES string tokens autoregressively.

### Loss Function

From the point of probabilistic modelling, the goal of VAE^26^ is to learn distribution *p*(*x*) where *x* corresponds to our data. However, when such distribution is complex, we can approximate it by using latent variable model *p*(*z*), which is assumed to be a standard Gaussian distribution, and *p*(*x*|*z*). The loss function *L* in the model plays a crucial role in guiding the training process and includes two primary components:

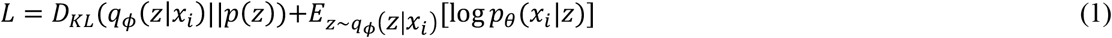

The first term in **Eq.1** and **Eq.4** is Kullback-Leibler (KL) divergence loss^27^, which measures the distance between two distributions. *q*_*ϕ*_ can be thought of as the encoder and *q*_*ϕ*_(*z*|*x*_*i*_) represents the process that the encoder encodes the data *x*_*i*_ into a latent variable *z*. The KL divergence term encourages the model to encode features of the data close to the prior distribution *p*(*z*) which is assumed to be a standard Gaussian distribution.

The second term in **Eq.1** is the reconstruction loss, where *p*_*θ*_ can be thought as a process of decoder trying to reconstruct the input. *Z* is sampled from *q*_*ϕ*_(*z*|*x*_*i*_) (**Eq.2**), and *p*_*θ*_(*x*_*i*_|*z*) is the process where our decoder receives the latent variable *z* and reconstructing the input from that. We implemented the second term in PyTorch, using negative log likelihood for a comparison of the generated output probabilities of SMILES tokens with the input SMILES. However, as *z* is sampled stochastically from a distribution, during backpropagation gradients would have to flow through this stochastic node, which is a non-differentiable function.

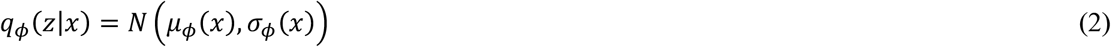

To overcome this, we apply reparameterization and instead of sampling *z* from *q*_*ϕ*_(*z*|*x*), we can sample *ε* from a normal distribution with a mean of zero and a variance of one and construct *z* from it as in **Eq.3**.

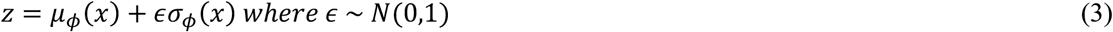

This approach helps retain the stochasticity of the distribution yet allows the gradient to flow through two deterministic functions *μ*_*ϕ*_ and *σ*_*ϕ*_. The updated loss functions *L* is then taking the form as in **Eq.4**.

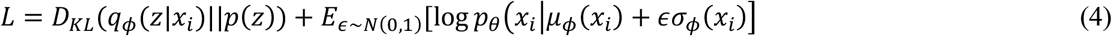

One of the challenges with training VAE is posterior collapse, which happens when the model can only generate limited samples and fails to accurately reconstruct or meaningfully represent the original input data. Moreover, posterior collapse can be worsened due to insufficient diversity of the dataset. In our case, VAE may overfit common patterns, particularly evident in the prevalence of carbon atoms observed in the SMILES token frequency distribution within our training data and across most molecules. Introducing a regularization term *β* has been shown to mitigate the issue of posterior collapse^28^. By default, the model aims to minimize both the data reconstruction loss and KL divergence at the same time. The hyperparameter *β* adjusts the emphasis on the KL divergence loss within the VAE objective function (**Eq.5**), encouraging diverse and informative latent representations and preventing the encoder from collapsing all data points into the same latent code.

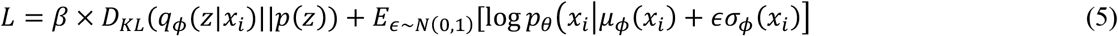

We assess the performance of TGVAE using two *β* scheduling strategies: monotonic and cyclical^29^. The former gradually increases *β* from low to high during training, while the latter periodically varies *β* between its lowest and highest values.

## Training and Parameter Settings

The training and parameter settings are carefully chosen to balance between model complexity and generalization. Specifically, we employ a stack of eight encoder and decoder blocks, each with a hidden dimension of 512 and a feed-forward dimension of 1024. To manage the dimensionality of the GAT within the encoder, which yields an output dimension of *hidden*_*dimension* * *num*_*heads*, we employ a single-head attention mechanism to mitigate potential overfitting arising from excessive dimensionality increase. In contrast, the decoder’s attention layer utilized a 16-head attention mechanism, aligning with the original attention layer’s dimension division for each head. During training, we employ the Adam optimizer^30^ with a learning rate set to 3e-4 and a weight decay of 1e-6. Our baseline model adopted a monotonic *β* schedule with weights ranging from 0.00005 to 0.0001. The model is trained for 100 epochs to ensure stable evaluation and robustness of results.

### Evaluation

Evaluation of the molecules generated by our TGVAE model adheres to the metrics outlined in the MOSES benchmark. In line with MOSES and the aim of exploring chemical space and generating diverse molecule set, we primarily focus on five key metrics: validity, uniqueness, novelty, internal diversity, and scaffold similarity. The benchmark suggests generating G set of 30,000 molecules. All metrics are calculated for the pool of valid molecules found in the G set. Each metric depends on G set and the test set R (reference set). Definitions of the metrics below are adapted from the MOSES benchmark.

Validity of molecules defined in **Eq.6** is evaluated using RDKit^31^ molecular structure analyzer that checks atoms’ valency and consistency of bonds in aromatic rings. Validity measures how well the model captures explicit chemical constraints such as proper valence:

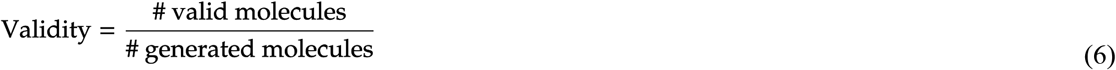

Uniqueness (**Eq.7**) counts the number of molecules that are unique or different from others in the G set. I.e., it checks that the model maintains diversity and does not converge solely on producing a few typical molecules. Here, we compute Unique@K for the first K = 10,000 valid molecules in the G set. If the number of valid molecules is less than K, we compute uniqueness on all valid molecules:

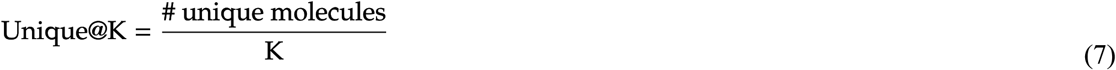

Novelty (**Eq.8**) measures the proportion of generated molecules absent in the training set. We prioritize this metric since we want to maximize the number of novel molecules. A low novelty score signals potential overfitting issues which we believe is the problem of most string-based models:

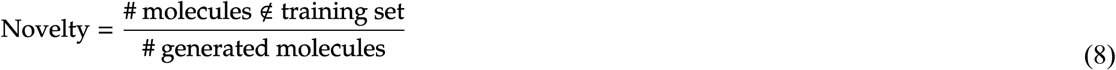

Internal diversity (IntDiv, **Eq.9**) evaluates the chemical variety within the generated molecule set, detecting frequent cases of posterior collapse in generative model. Low internal diversity scores, which are associated with posterior collapse, indicate the model generates a limited range of samples, neglecting certain areas of the chemical space. IntDiv ranges between 0 and 1:

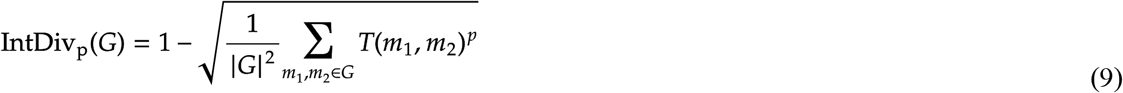

Scaffold similarity (**Eq.10**) compares frequencies of Bemis-Murcko scaffolds containing ring structures and linker fragments connecting these rings. Using *c*_s_(*A*) to denote the frequency of scaffolds in molecules from a placeholder set A, and S as a set of molecules appearing in either G or R, the metric is defined as cosine similarity. This metric reveals the similarity between scaffolds present in the generated and reference datasets. A lower scaffold similarity may indicate infrequent production of certain scaffolds from the reference set. Notably, since the scaffold test set contains scaffolds absent in both the train and test sets, higher scaffold similarity with this set suggests the model’s capability to generate novel scaffolds not present in the training set:

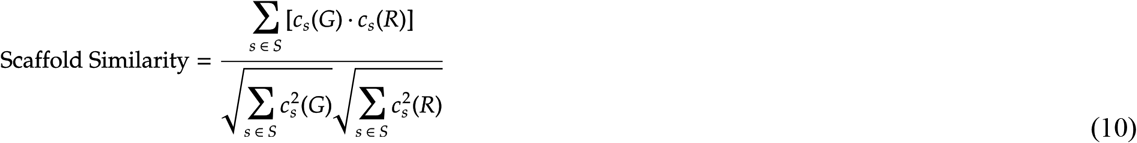

## RESULTS

### Model validation

We compare TGVAE model’s performance with other string-based models. The generated molecules are evaluated by five metrics as described above: validity, uniqueness, novelty, internal diversity, and scaffold similarity to the scaffold test set. While achieving a comparable level of validity and uniqueness with other models, TGVAE outperforms other models in generating novel molecules, novel scaffolds, and more diverse set (**Table 1**). As shown in **Table 1**, IntDiv1 and IntDiv2 (for p=1 and p=2 in Eq.9, respectively) for the training set are 0.857 and 0.851, respectively. All string-based model can only achieve lower or equal results in comparison to the training set. TGVAE outperformed these models, including internal diversity of the molecules, which is higher than the training set. We tested training of the model with two types of *β* schedule: monotonic (M) and cyclical (C) with one attention head (H) in the GAT layer (1H/M, 1H/2C, 1H/4C, 1H/8C, **Table 1**). Results showed that with a small decrease in validity, cyclical schedule helps the model generate higher fraction of novel molecules. We also investigated the consequences of increasing the number of attention heads in the GAT layer (1H/M, 2H/M, 4H/M, 8H/M, **Table 1**). This approach led to increased novelty, higher fraction of novel scaffolds yet some tradeoff in validity of generated molecules was observed. In general, TGVAE showed improved performance in terms of novelty and chemical variety compared to previous string-based models. Examples of the molecules generated by TGVAE are illustrated in **Figure 2**.

TGVAE maintained adherence to properties of the MOSES original distribution. Results demonstrate that molecules generated by TGVAE exhibit a slightly broader distribution of MW (**Figure 3A**), sampling both smaller and larger molecules relative to the test set. Despite this variability, the distributions of key molecular properties such as logP, QED (quantitative estimate of drug-likeness), and SA (synthetic accessibility) (**Figure 3B-D**) in the generated set closely resembles those observed in the test set (**Figure 3**, in brackets - Wasserstein^32^ distance from TGVAE to MOSES test set). This alignment underscores the capability of TGVAE to generate diverse and novel molecules that maintain critical physicochemical characteristics, thereby validating its efficacy in diverse chemical design applications.

**Figure 2.**
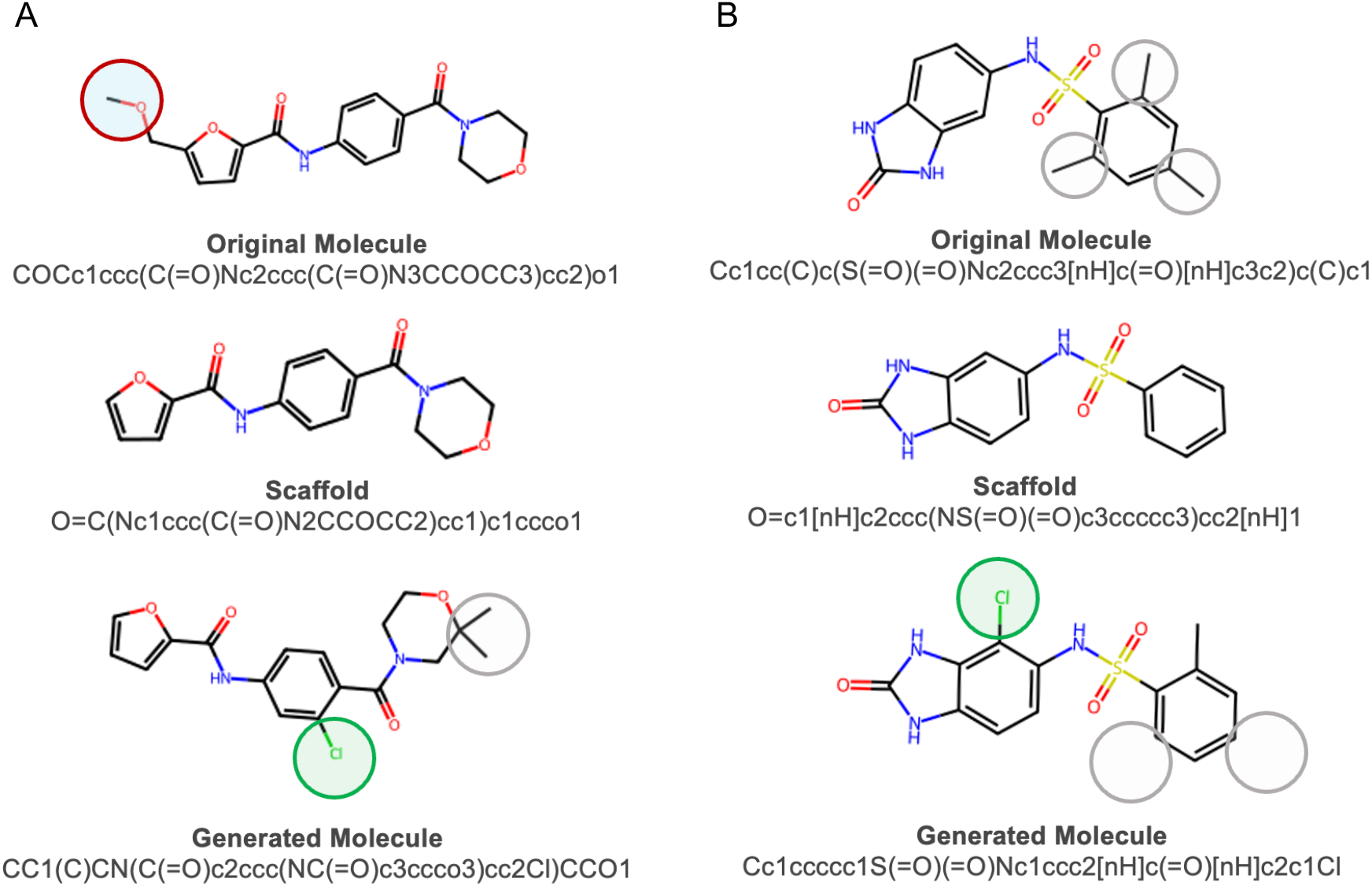
Two examples (A and B) of molecules generated by TGVAE. Starting from the original molecule (top row), the scaffold is generated (middle row) and decorated with the new functional groups (bottom row).

**Table 1.**
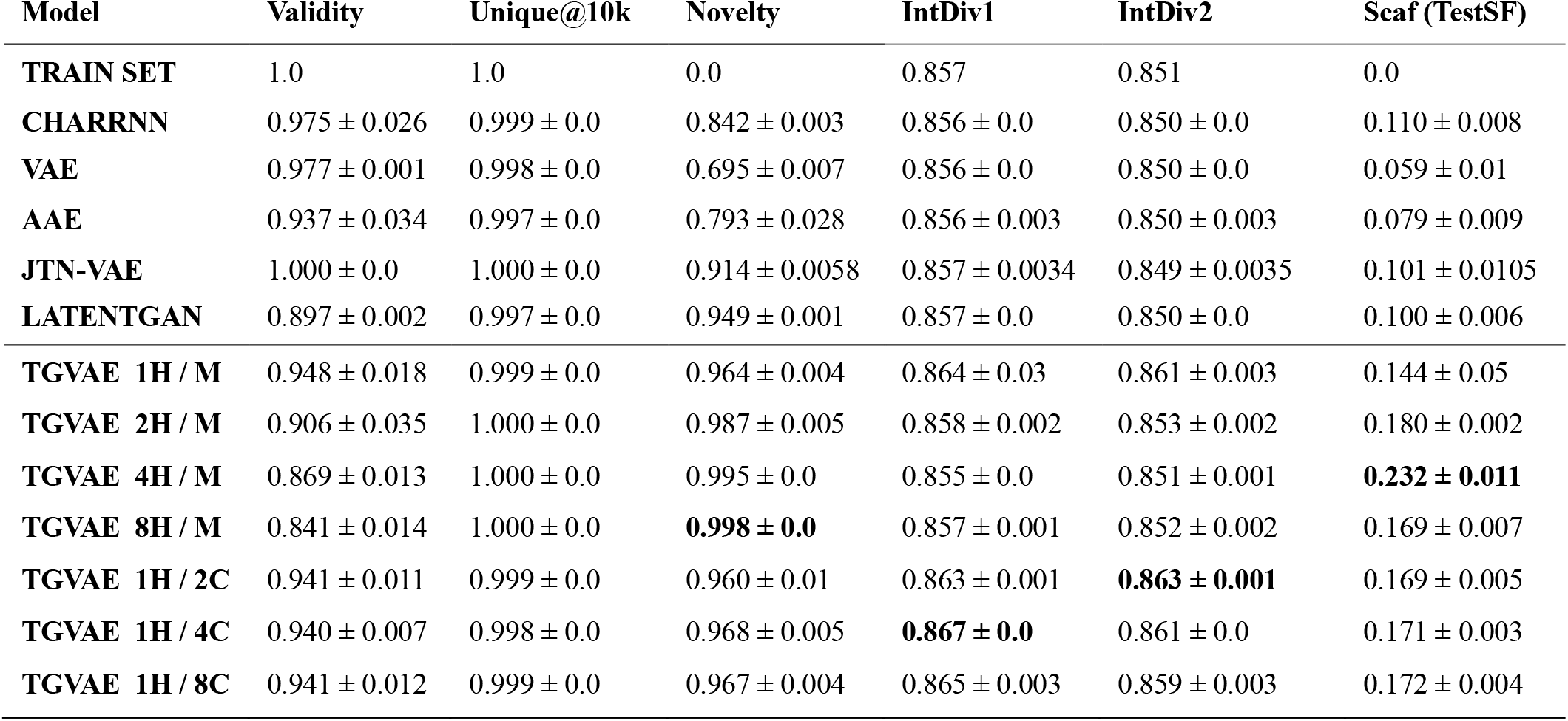
MOSES performance metrics for the baseline models and various TGAVE setups. Fraction of valid molecules, fraction of unique molecules from 10k set, novelty of molecules, internal diversities 1 and 2, and scaffold similarity test. Standard deviations are calculated from three independent runs.

| Model | Validity | Unique@10k | Novelty | IntDiv1 | IntDiv2 | Scaf (TestSF) |
| --- | --- | --- | --- | --- | --- | --- |
| TRAIN SET | 1.0 | 1.0 | 0.0 | 0.857 | 0.851 | 0.0 |
| CHARRNN | 0.975 ± 0.026 | 0.999 ± 0.0 | 0.842 ± 0.003 | 0.856 ± 0.0 | 0.850 ± 0.0 | 0.110 ± 0.008 |
| VAE | 0.977 ± 0.001 | 0.998 ± 0.0 | 0.695 ± 0.007 | 0.856 ± 0.0 | 0.850 ± 0.0 | 0.059 ± 0.01 |
| AAE | 0.937 ± 0.034 | 0.997 ± 0.0 | 0.793 ± 0.028 | 0.856 ± 0.003 | 0.850 ± 0.003 | 0.079 ± 0.009 |
| JTN-VAE | 1.000 ± 0.0 | 1.000 ± 0.0 | 0.914 ± 0.0058 | 0.857 ± 0.0034 | 0.849 ± 0.0035 | 0.101 ± 0.0105 |
| LATENTGAN | 0.897 ± 0.002 | 0.997 ± 0.0 | 0.949 ± 0.001 | 0.857 ± 0.0 | 0.850 ± 0.0 | 0.100 ± 0.006 |
| TGVAE 1H / M | 0.948 ± 0.018 | 0.999 ± 0.0 | 0.964 ± 0.004 | 0.864 ± 0.03 | 0.861 ± 0.003 | 0.144 ± 0.05 |
| TGVAE 2H / M | 0.906 ± 0.035 | 1.000 ± 0.0 | 0.987 ± 0.005 | 0.858 ± 0.002 | 0.853 ± 0.002 | 0.180 ± 0.002 |
| TGVAE 4H / M | 0.869 ± 0.013 | 1.000 ± 0.0 | 0.995 ± 0.0 | 0.855 ± 0.0 | 0.851 ± 0.001 | **0.232 ± 0.011** |
| TGVAE 8H / M | 0.841 ± 0.014 | 1.000 ± 0.0 | **0.998 ± 0.0** | 0.857 ± 0.001 | 0.852 ± 0.002 | 0.169 ± 0.007 |
| TGVAE 1H / 2C | 0.941 ± 0.011 | 0.999 ± 0.0 | 0.960 ± 0.01 | 0.863 ± 0.001 | **0.863 ± 0.001** | 0.169 ± 0.005 |
| TGVAE 1H / 4C | 0.940 ± 0.007 | 0.998 ± 0.0 | 0.968 ± 0.005 | **0.867 ± 0.0** | 0.861 ± 0.0 | 0.171 ± 0.003 |
| TGVAE 1H / 8C | 0.941 ± 0.012 | 0.999 ± 0.0 | 0.967 ± 0.004 | 0.865 ± 0.003 | 0.859 ± 0.003 | 0.172 ± 0.004 |

**Figure 3.**
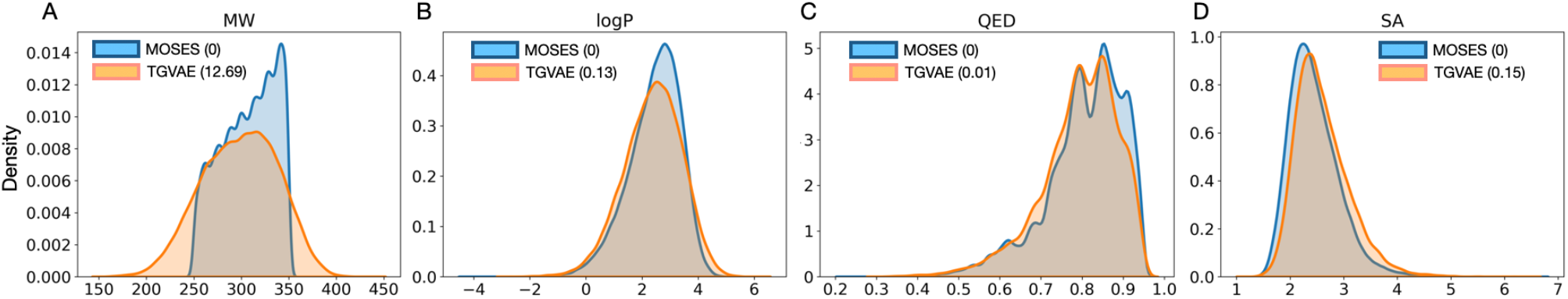
Distribution of chemical properties of MOSES test set and molecules generated by TGVAE. In brackets is Wasserstein distance to MOSES test set. (A) Molecular weight (MW in g/mol). (B) Octanol-water partition coefficient (logP), (C)Quanqtitative estimation of drug-likeness (QED). (D) Synthetic accessibility score (SA). Results from one of the three simulations.

### Novel structure generation

By training on the datasets including 1.5M molecules, we further compared the generated by TGVAE model structures with those in PubChem^33^. PubChem is a chemical information resource and database available from National Institutes of Health and provides information about chemical structures, identifiers, chemical and physical properties, biological activities, and more data on small and larger molecules such as nucleotides, peptides, lipids, carbohydrates, and other chemically modified macromolecules.

To begin the comparison of TGVAE-generated scaffolds with the catalog of the available at PubChem molecules (118M), we utilized PUG-REST^34^ – a Representational State Transfer (REST) PubChem web service that allows access to and provides information on the available records. PUG-REST handles request and performs the search by using e.g., structure identify such as SMILES string format input, and is capable to return various compound properties. Using this functionality of PubChem server, we implemented in-house python scripts capable of systematic comparison of each TGVAE-generated scaffold against every record within the PubChem repository of chemical compounds. For the scaffold evaluation, we defined the search to return the information about the structure similarity to existing molecules and the count of substructures associated with a given molecule.

In PubChem, the structure similarity is done by 2D Tanimoto coefficient^35,36^ calculation while the substructure identification refers to finding a specific part of a molecule that can be found within known larger molecules. By identifying molecules, which are not a substructure to any known molecule in parallel with evaluation of a lack of similarity, we determined the presence of novel molecules. Conversely, a superstructure analysis, i.e., analysis if is a given structure contains the entire smaller molecule as a part of its own composition, helps in understanding how the known pharmacophores or functional groups fit within the generated molecules, indicating potential biological activity. We thus present the similarity, substructure, and superstructure count comparative analyses with PubChem database as elucidating the novelty and potential utility of the TGVAE-generated molecules. In addition, this approach helped ensure that the structures either represent entirely new chemical entities or novel variations of existing compounds (**Figure 4**).

**Figure 4.**
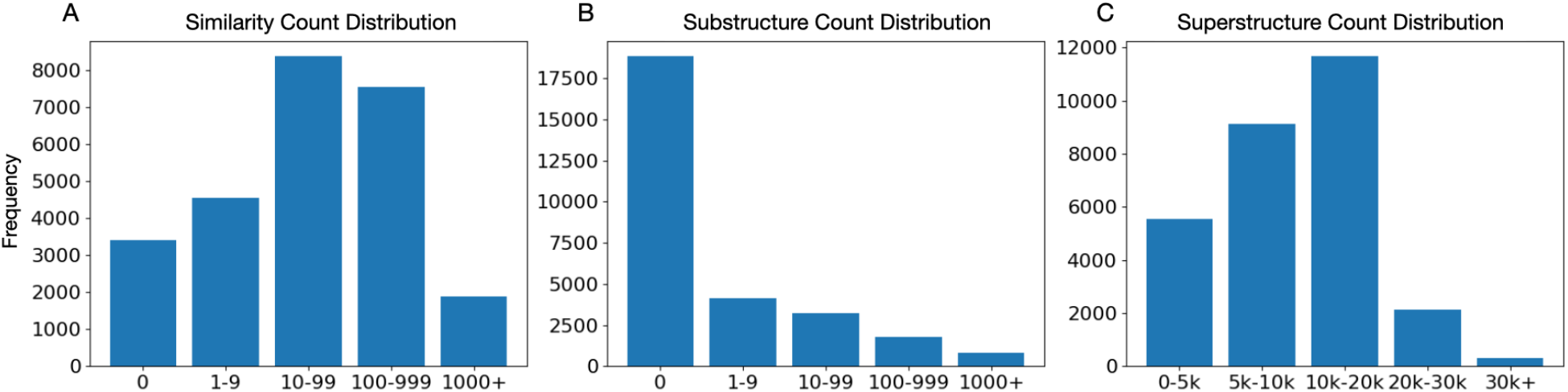
Distributions extracted from PubChem for generated valid molecules. (A) Similarity count distribution. (B) Substructure count distribution. (C) Superstructure count distribution. On the plots, x axes represent bins with the count of PubChem structures similar (A), substructures (B), superstructures (C) to those generated by TGVAE; y axis – frequency of TGVAE generated scaffolds in each of the bins.

From the pool of valid molecules generated by the TGVAE model, over 3k molecules showed zero similarity with any molecule present in the PubChem database; over 4k molecules had up to 10 similar structures found; while over 8k, 7k, and nearly 2k of the generated structures had over 10, 100, and 1000 similar structures at PubChem, respectively (**Figure 4A**). Over 50% structures generated by TGVAE were not substructures to any from PubChem (**Figure 4B**) while all the generated structures were superstructures to smaller fragments (**Figure 4C**). The latter observation is important in evaluation of the probability of the TGVAE novel molecules to be synthesized and possessing biological activity.

### Evaluation of the novel structures

We further compared the >3k novel structures with zero similarity with the PubChem database (*zero-similarity structures* henceforth, **Figure 4A**) to the 30k generated molecules and the MOSES test set. We focused on several properties related to lipophilicity and physicochemical character, including logP, HBD (hydrogen bond donors), HBA (hydrogen bond acceptors), QED, NH (amine) and OH (hydroxyl) groups, as well as number of heavy atoms (**Figure 5A-F**). We noticed that these properties differed for the structures with zero-similarity and indicated slightly lower hydrophobicity than the other two sets (30k, MOSES) with the mode of logP distribution shifting from above to less than two (**Figure 5A**). In general, compounds with a logP around that value are desirable in drug design because of their good permeability across cell membranes while maintaining adequate solubility in aqueous environments (such as blood plasma). The observed lower HBD trend (**Figure 5B**) among the zero-similarity group suggested reduced solubility in water due to potentially reduced hydrogen bonding with water molecules. A higher number of HBA on the other hand, (**Figure 5C**) suggested some degree of solubility. These changes in HBD and HBA trends suggested that TGVAE model generates novel molecules with hydrogen bonding properties maintaining a balance between enhanced cell membrane permeability and increased interactions with polar environments.

Although the observed slight shift toward the lower values in QED distribution among the zero-similarity structures might suggest less favorable range of drug-likeness (**Figure 5D**), a QED mode of ∼0.7 suggests TGVAE generates highly potent candidates for fragment-based drug design and prospective optimization. Generated compounds with this range might have other advantages, such as strong efficacy or selectivity.

Finally, a reduced number of NH (amine) and OH (hydroxyl) groups as well as reduced number of heavy atoms observed in the generated structures with zero-similarity (**Figure 5E-F**) could significantly influence their drug-likeness and pharmacokinetic properties. I.e., the pharmacokinetic properties could be improved since fewer heavy atoms may lead to smaller or more compact molecules. On the other hand, reducing the number of heavy atoms might negatively impact molecule’s structural complexity and ability to interact effectively with its biological target. This however suggests that the zero-similarity generated molecules represent potent candidates to optimization approaches and such smaller essential fragments can be further modified to achieve enhanced drug-like properties.

**Figure 5.**
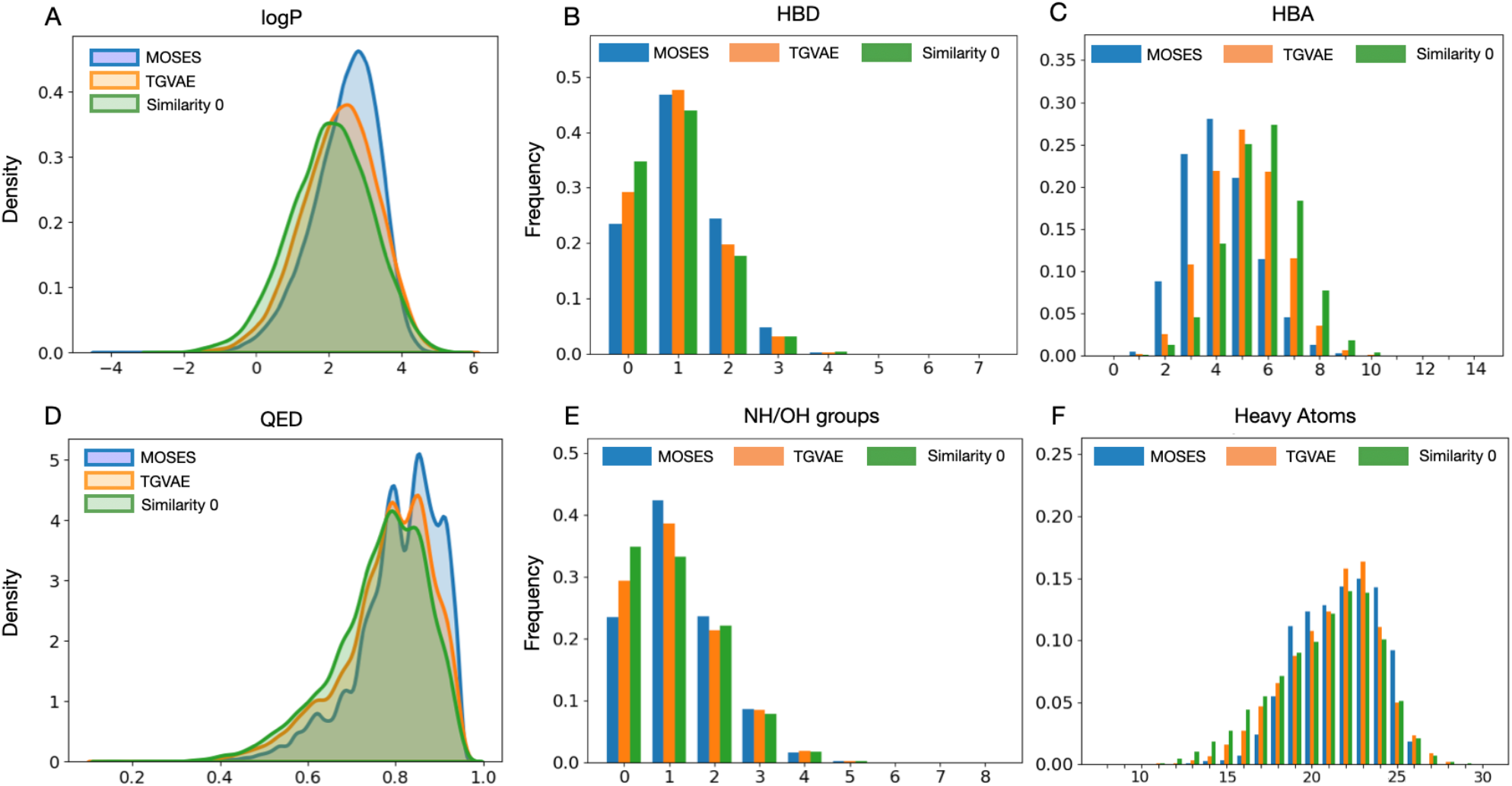
Density distributions for the sets of structures with similarity zero, 30k generated molecules, and MOSES set. (A)logP. (B) HBD. (C) HBA.(D) QED. (E) Number of NH/OH groups. (F) Number of heavy atoms. Calculated using all three simulations.

Code availability: https://github.com/nnhoangtrieu/TGVAE

## CONCLUSIONS

Traditional drug discovery methods often rely on existing chemical libraries and known compounds, which can limit the exploration of new chemical spaces. Generative AI emerges as a powerful tool in the field of drug discovery, particularly in the generation of novel structures or drug-like molecules with improved properties. These properties include enhanced binding affinity, improved pharmacokinetics, reduced off-target effects, and lower toxicity profiles. The development, implementation, and application of the TGVAE model discussed in this article, capable of novel structures generation, facilitates identification of the lead compounds with high therapeutic potential and provides opportunity to accelerate drug design and new drug discovery, Our approach not only complements traditional medicinal chemistry approaches by uncovering unique chemical entities but also provides a robust framework for addressing many diseases that have been challenging to treat using conventional drug discovery methods. The innovative capacity of TGVAE to navigate and exploit previously unexplored chemical spaces signifies a transformative advancement in AI. Collaborative efforts between interdisciplinary teams comprising chemists, biologists, data scientists, and clinicians are however necessary in expanding the full potential of AI to drug development.

## ACKNOWLEDGEMENTS

All computations were performed using HPC and DGX computing clusters at H. Lee Moffitt Cancer Center and Research Institute.

## Abbreviations

AI: Artificial Intelligence
GAT: Graph Attention Network
GNN: Graph Neural Network
KL: Kullback-Leibler
VAE: Variational Autoencoder
TGVAE: Transformer Graph Variational Autoencoder
MOSES: Molecular Sets
SMILES: Simplified Molecular Input Line Entry System

